# A phased chromosome-level genome resource for a myrtle rust susceptible *Syzygium luehmannii*

**DOI:** 10.1101/2025.05.12.653576

**Authors:** Peri A. Tobias, Jacob Downs, Alyssa M. Martino, Richard J. Edwards

## Abstract

*Syzygium luehmannii* is an Australian east coast endemic tree within the family Myrtaceae. *Syzygium luehmannii* is not known to be highly susceptible to the parasitic fungus causing myrtle rust, *Austropuccinia psidii*, however infections have been reported in the field, and in controlled inoculations. The capacity for this exotic pathogen to parasitise host trees, suggest that molecular targets are present in susceptible plants. While understanding resistance phenotypes is important for tree breeding and management, determining the key drivers for susceptibility may also provide useful additional research targets to avert infection. While there are several genome resources for plants within the large and globally diverse *Syzygium* genus, there is no diploid genome assembly (2n = 22), and no genome for *S. luehmannii*. We assembled the genome for *S. luehmannii* into the pseudo-phased, 11 chromosome pairs here termed haplotype A (370 Mbp) and B (357 Mbp). We annotated the predicted protein coding genes, and we specifically annotated the *nucleotide-binding leucine rich repeat* (NLR) type resistance genes as a useful resource for plant:pathogen studies. The high quality of this genome provides a base for studies on myrtle rust resistant and susceptible hosts to understand mechanisms of infection.

## Background

*Syzygium luehmannii* (F.Muell.) L.A.S. Johnson is an Australian east coast endemic tree within the family Myrtaceae. The natural distribution for this sub-tropical rainforest species is the coastal regions north of Kempsey, NSW (Royal Botanic Gardens and Domain Trust) with disjunct populations endemic to Cape York peninsula and northeast Queensland (CSIRO 2020). Unlike some other species within the *Syzygium* genus, such as *S. maire* (A. Gunn.) Sykes et Garn.-Jones and *S. jambos* L. (Alston), *S. luehmannii* is not known to be highly susceptible to the parasitic fungus causing myrtle rust, *Austropuccinia psidii* (Beenken 2017). Observed infections have, however, been reported in the field and in controlled inoculations in Australia (Morin et al. 2012; Soewarto et al. 2019; Tobias et al. 2018). Previous research identified 25 percent of plants as highly susceptible and 75 percent resistant, with 30 percent presenting classic hypersensitive response (HR), in a controlled inoculation with the pandemic strain of *A. psidii* (Tobias et al. 2018). These data suggest that a potential pathogen target is present in susceptible plants for the pathogen to successfully parasitise its host, and conversely a specific and rapid genetic response in resistant plants. While understanding resistance phenotypes is important for tree breeding and management, determining the key drivers for susceptibility may also provide useful additional research targets to avert infection (Henningsen et al. 2021), or to explain mechanisms of infection (Ceulemans 2021-effector hubs). While there are several genome resources for plants within the large and globally diverse *Syzygium* genus (Low et al. 2022; Ouadi et al. 2022; Balkwill et al. 2024), there is no diploid genome assembly (2n = 22), and no genome for *S. luehmannii*. We assembled the genome for *S. luehmannii* into the pseudo-phased, diploid chromosome pairs providing a useful resource for studies into the mechanisms underlying susceptibility to myrtle rust infection.

## Results and Discussion

Here, we present a pseudo-phased, chromosome level genome for *S. luehmannii*. The genome completeness, using Benchmarking Universal Single Copy conserved Orthologs (BUSCOs) (Manni et al. 2021), is 98.4% and 98.6% for each haplotype respectively (**Table 1**). We annotated the predicted protein coding genes, and we specifically annotated the *nucleotide-binding leucine rich repeat* (NLR) type resistance genes for each haplotype. As previously determined in the genome data from diverse Myrtaceae species, the NLR-type predicted genes are highly clustered with an expanded TNL-type class of genes (Thrimawithana et al. 2019; Balkwill et al. 2024; Christie et al. 2016; Chen et al. 2023). For example, the chromosome-level but collapsed haploid genome for *Syzygium maire* (Balkwill et al. 2024) was found to have 642 NLR-type genes and of these 333 were TNL-type. Similarly, the *S. luehmannii* genomes had 655 (633) NLRs and 328 (327) TNLs for haplotype A and B respectively, showing high conservation of numbers using the same FindPlantNLRs software pipeline (Chen et al. 2023). We determined the presence of non-conventional NLRs containing jacalin domains (PF01419), potentially replacing the LRR role, as also determined for *S. maire* (Balkwill et al. 2024). These duplicated jacalin domain containing NBARC predicted genes are an apparent common feature in the Myrtaceae, presenting an interesting research opportunity for investigation (Esch & Schaffrath 2017). Our research presents the allelic variance within paired chromosomes in a single diploid tree. The high quality of the scaffolded genome and annotation of haplotype chromosome pairs provides a useful resource for studies into the mechanisms underlying susceptibility to myrtle rust infection as well as comparative genomic analyses within the Myrtaceae family.

**Table 1.**
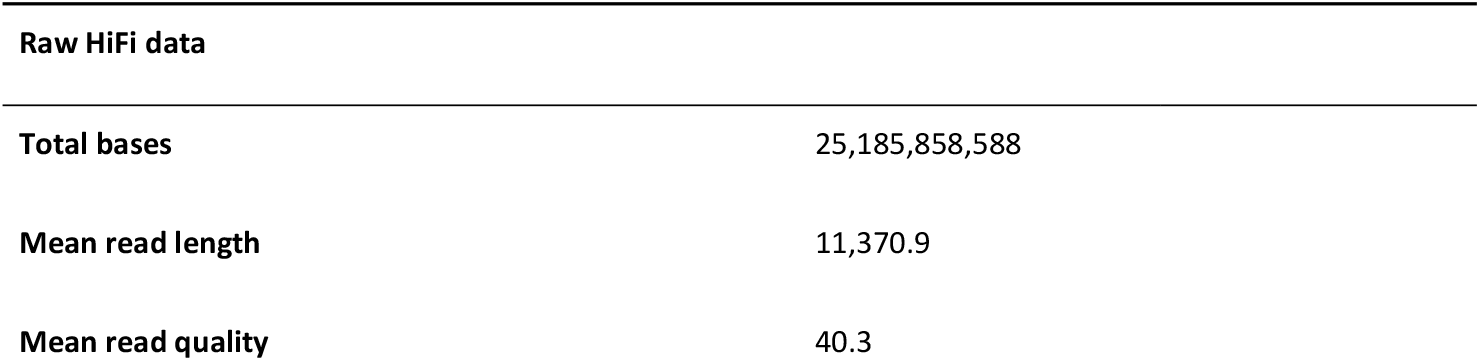
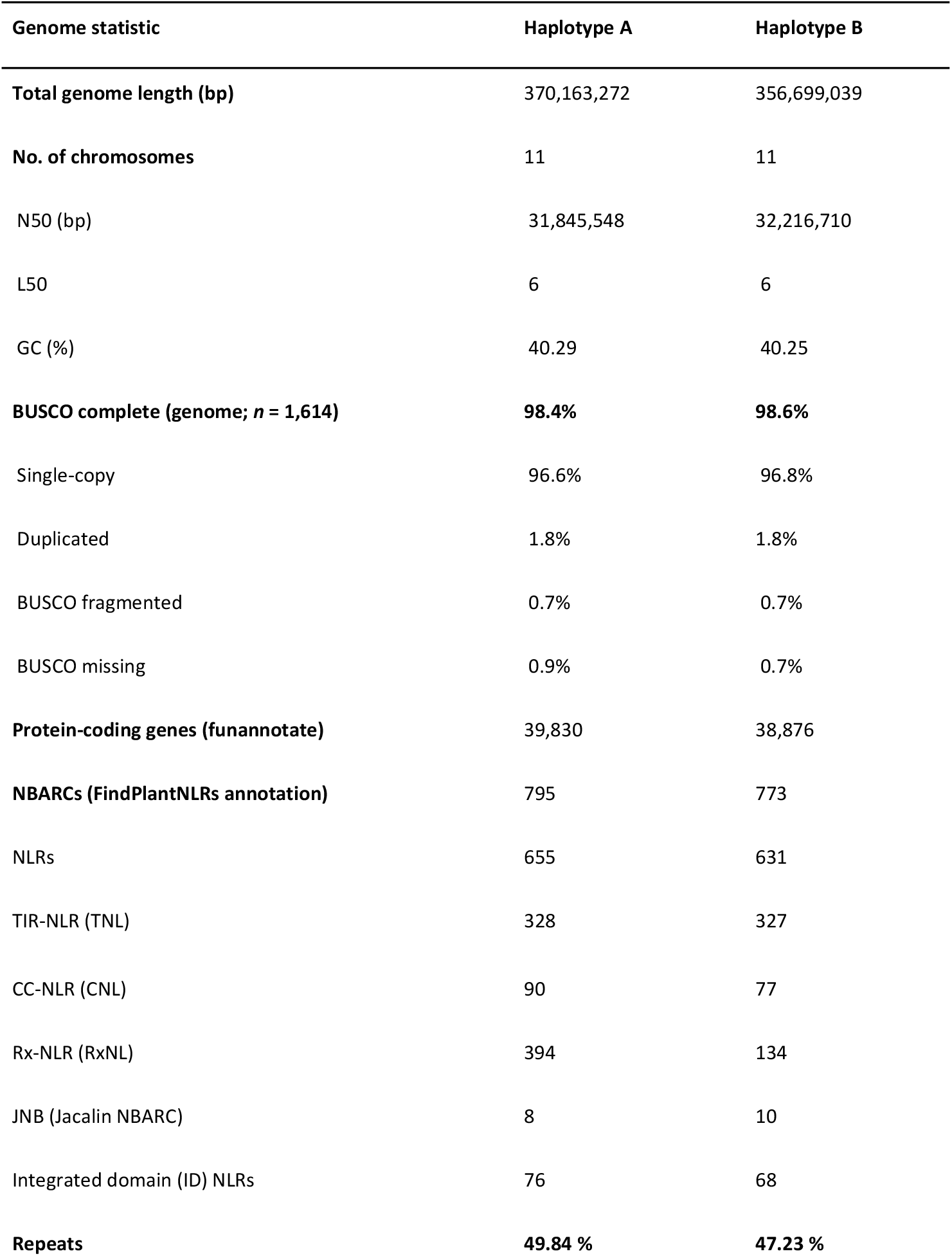
Raw data statistics from NanoPlot (De Coster & Rademakers 2023) and genome statistics for both sets of *Syzygium luehmannii* chromosomes using Quast and BUSCOs (Simão et al. 2015; Gurevich et al. 2013). Annotation statistics based on FindPlantNLRs and Funannotate (Chen et al. 2023; Palmer & Stajich 2023). *Nucleotide binding site leucine rich repeat* (NLR) type resistance genes were predicted, annotated and classified including those incorporating integrated domains (ID).

| Raw HiFi data |  |
| --- | --- |
| Total bases | 25,185,858,588 |
| Mean read length | 11,370.9 |
| Mean read quality | 40.3 |

| Genome statistic | Haplotype A | Haplotype B |
| --- | --- | --- |
| **Total genome length (bp)** | 370,163,272 | 356,699,039 |
| **No. of chromosomes** | 11 | 11 |
| N50 (bp) | 31,845,548 | 32,216,710 |
| L50 | 6 | 6 |
| GC (%) | 40.29 | 40.25 |
| **BUSCO complete (genome; *n* = 1,614)** | **98.4%** | **98.6%** |
| Single-copy | 96.6% | 96.8% |
| Duplicated | 1.8% | 1.8% |
| BUSCO fragmented | 0.7% | 0.7% |
| BUSCO missing | 0.9% | 0.7% |
| **Protein-coding genes (funannotate)** | 39,830 | 38,876 |
| **NBARCs (FindPlantNLRs annotation)** | 795 | 773 |
| NLRs | 655 | 631 |
| TIR-NLR (TNL) | 328 | 327 |
| CC-NLR (CNL) | 90 | 77 |
| Rx-NLR (RxNL) | 394 | 134 |
| JNB (Jacalin NBARC) | 8 | 10 |
| Integrated domain (ID) NLRs | 76 | 68 |
| **Repeats** | **49.84 %** | **47.23 %** |

**Figure 1.**
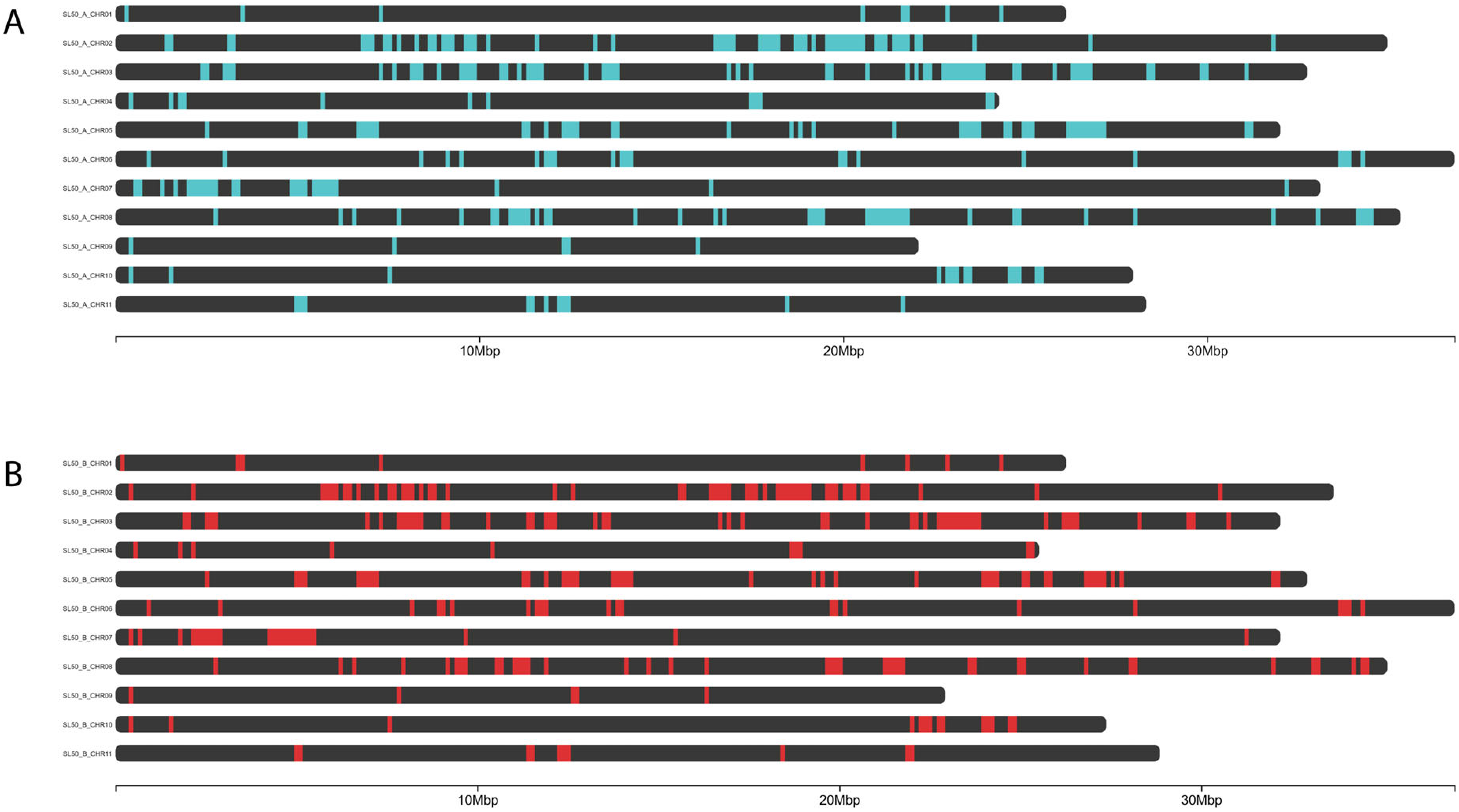
Genomic clustering of predicted *nucleotide binding leucine rich repea*t genes (NLRs) on the diploid chromosomes of *Syzygium luehmannii*. NLR clusters are visually presented using chromomap (cite) indicating variation in cluster size and locations between chromosome pairs. Haplotypes A (A) B (B) NLRs are represented with blue and red segments respectively.

## Methods

### Sample collection

*Syzygium luehmannii* tubestock plants (50) were commercially sourced from Burringbar Rainforest Nursery, New South Wales (NSW) in August 2018 and labelled SL1 – SL50. Seed for these plants was sourced from parent trees located at Wooyung and Bilinudgel on the NSW north coast. The plants were inoculated in October 2018 with the pandemic strain of *Austropuccinia psidii* (Au-3 glasshouse increased 622) at the Plant Breeding Institute (PBI), University of Sydney, using methods described previously (Tobias et al. 2018). Ratios were consistent with an earlier experiment (Tobias et al. 2018) at 74% resistant (score 1-3) and (26%) susceptible (score 4-5). One highly susceptible plant, designated SL50, was planted and cuttings were made for additional controlled inoculations. Controlled inoculations were repeated on the established cuttings in November 2021 and in June 2022. Phenotypes were consistent across every controlled inoculation, with heavy infection on susceptible plants. A further controlled inoculation was done in February 2023 confirming the repeated susceptible phenotype. Notably, on the 21 October 2022, myrtle rust field infection was recorded for the first time on the planted specimen. The field infection was notable in that it confirmed the highly susceptible status of SL50. The plant was selected for genome assembly and annotation, as an example of a highly myrtle rust-susceptible individual and a specimen was vouchered in December 2022 at the National Herbarium of NSW, Royal Botanic Gardens, Sydney (NSW 1124137).

### DNA extraction and sequencing

High Molecular Weight DNA was extracted using a protocol adapted from the sorbitol wash method developed by Jones et al. (2021), and the HMW plant DNA CTAB extraction method developed by Hilario (2018). Small (1 cm^2^) pieces of the leaf lamina were placed into 1.5 mL centrifuge tubes with a small amount (a few mg) of PVP40000. The tissue was flash frozen in liquid nitrogen and ground into a powder with a micropestle while frozen. Two rounds of sorbitol wash were used on the samples, in accordance with the protocol (Jones et al., 2021), until the supernatant was no longer viscous. Then DNA was extracted, according to the protocol (Hilario, 2018), using CTAB buffer with an incubation step at 56 °C for 2 hours, in a water bath. Apart from an extended RNase A incubation step of 10 mins, the protocol was followed until the precipitation step. The DNA formed a visible gelatinous precipitate at this stage and was left to continue precipitating at -20 °C overnight. The following day the DNA was collected by centrifugation at 3000 rpm for 25 min at room temperature and cleanup steps followed according to the protocol before dissolving in 10 mM Tris-HCL pH 8.0. The DNA was quantified, and purity was checked using a NanoDrop™ 2000c spectrophotometer and a QuBit™ 2.0 Fluorometer. Purified HMW DNA (10 × 1.5 µL tubes) was couriered under an Australian Department of Agriculture, Water, and the Environment import permit to the Australian Genome Research Facility (AGRF), Brisbane, Australia, for cleanup, size selection, library prep and sequencing on Pacific Biosciences of California, Inc. (PacBio) Sequel II.

### Formaldehyde cross-linking of leaves for HiC sequencing

We prepared samples according to the Phase Genomics Proximo Hi-C Fungal Kit protocols for chromatin crosslinking, followed by library preparation and Illumina MiSeq sequencing at Ramaciotti Centre for Genomics, University of NSW, Australia. Five grams of leaf material was immersed in 2% (v/v) sodium hypochlorite for 2 mins for surface sterilisation and then washed with ultrapure water. The leaves were cut into strips and immersed in 1% (v/v) formaldehyde for 40 minutes with intermittent mixing. The crosslinking reaction was quenched with the addition of glycine to a final concentration of 125 mM for 15 minutes. The crosslinked leaves were then ground to a powder after freezing with liquid nitrogen. As the samples were not washed prior to grinding, a wash step was done prior to lysis during library preparation.

### Data processing, genome assembly and curation

Details of all the software, versions, parameters and scripts used to process the data are available here on github (Tobias 2025). In brief, the PacBio Sequel II ccs.bam file (17G) was downloaded 4 March 2023 and filtered of adaptors using HiFiAdapterFilt. We ran NanoPlot on the filtered data (2,214,940 total reads) to check statistics with mean raw read lengths and quality at 11,370.9 bases and 40.3 respectively. We received 47 Gigabytes of HiC paired-end fastq.gz data from the Ramiacotti Centre for Genomics, UNSW on the 27 April 2023. To filter pathogen (*A. psidii*) from host (*S. luehmannii*) sequences, we mapped the ccs data to the diploid *A. psidii* genome (Edwards et al. 2022)(https://zenodo.org/records/6476632) including the mitochondrial sequence, using Minimap2. We used the unmapped reads (98%) to build the genome and incorporated the HiC reads with Hifiasm. The diploid coverage for the plant was approximately 35 times. We scaffolded the genome with HiC data using Juicer (Neva C. Durand et al. 2016) and 3D-DNA (Dudchenko et al. 2017) pipelines using Juicebox (Neva C Durand et al. 2016) to visualize scaffolds for manual correction. We ran reference-based scaffold anchoring and super-scaffolding on each of our HiC haploid scaffolded genomes independently with pafscaff (Field et al. 2020), using the NCBI *Melaleuca quinquenervia* reference genome (GCA_030064385.1). We then concatenated the scaffolded and unplaced fasta files for a complete set of contigs for each haplotype and then ran Telociraptor (Edwards 2023) followed by Chromsyn on just the chromosome-level contigs to compare synteny between haplotype A and B. Initially the synteny plots indicated some errors in scaffolding, that were manually adjusted before rerunning Chromsyn. We then ran the genomes through contamination screening using FCS-GX (Astashyn et al. 2024) and Taxolotl (Tobias et al. 2022) and removed contigs that were flagged as non-plant or plastid to retain only genomic DNA sequences. The resulting principle and secondary pseudo-haplotype assemblies were labelled SL_A.v1.1.fasta and SL_B.v1.1.fasta.

### Annotation

We repeatmasked each haploid genome using Repeatmodeler (Smit & Hubley 2019) and Repeatmasker. Following this we mapped publicly available RNAseq data from NCBI: PRJNA356336 (Tobias et al. 2018) to the repeatmasked genomes independently with Hisat2 (Kim et al. 2019). The paired data was concatenated to make a single R1 and R2 fastq file, then trimmed of adaptors and low-quality reads using Fastp (Chen et al. 2018) in default mode and outputs processed with Samtools (Li et al. 2009) to make bam files. The mapped RNAseq bam files were used as evidence to predict protein coding genes within the Funannotate pipeline (Palmer & Stajich 2023). The protein annotations were run with interproscan5 (Jones et al. 2014). We ran both haplotypes through the FindPlantNLRs (Chen et al. 2023) docker installed pipeline to annotate the predicted resistance genes. The NLR gene locations were visualised with ChromoMap (Anand & Rodriguez Lopez 2022).

## Data availability

Raw data associated with this project is available here: https://www.ncbi.nlm.nih.gov/bioproject/PRJNA1099473. Diploid genome data is available at the following NCBI accessions: JBEWWB000000000, JBEWWC000000000. Whole genome annotation data is available here: https://doi.org/10.5281/zenodo.15321351 and NLR annotation data is available here: https://github.com/peritob/Syzygium-luehmannii.

## Acknowledgements

We thank the Australian Botanic Garden, Mt Annan, for use of the molecular facilities. We would like to acknowledge the contribution of the Plant Pathogen ‘Omics Initiative consortium in the generation of data used in this publication. The Initiative is supported by funding from Bioplatforms Australia, enabled by the Commonwealth Government National Collaborative Research Infrastructure Strategy (NCRIS).

## Abbreviations

BUSCO: Benchmarking Universal Single-Copy Orthologs
CC-NLR (CNL): Coiled-coil Nucleotide-binding Leucine-rich Repeat
CTAB: Cetyltrimethylammonium Bromide
HiC: Chromosome Conformation Capture
HMW: High Molecular Weight
ID: Integrated Domain
JNB: Jacalin NBARC
NBARC: Nucleotide-binding Apoptotic protease-activating factor-1, R protein, and CED-4 domain
NLR: Nucleotide-binding Leucine-rich Repeat
RxNL: CNL disease resistance protein that mediates resistance to potato virus X in *Solanum tuberosum*.
TIR-NLR (TNL): Toll/Interleukin-1 Receptor Nucleotide-binding Leucine-rich Repeat

## Author contributions

PT initiated the research, sourced funding and plants, inoculated/scored the plants with *A. psidii*, assembled, scaffolded and annotated the genome, submitted the data to NCBI and wrote the manuscript. JD extracted HMW DNA and cross-linked leaf samples. AM inoculated the plants for DNA extraction and ran NLR analysis. RE ran genome contamination screening. All authors contributed to editing the manuscript.

## Author notes

The authors declare no conflict of interest.

